# AnAms1.0: A high-quality chromosome-scale assembly of a domestic cat *Felis catus* of American Shorthair breed

**DOI:** 10.1101/2020.05.19.103788

**Authors:** Sachiko Isobe, Yuki Matsumoto, Claire Chung, Mika Sakamoto, Ting-Fung Chan, Hideki Hirakawa, Genki Ishihara, Hon-Ming Lam, Shinobu Nakayama, Shigemi Sasamoto, Yasuhiro Tanizawa, Akiko Watanabe, Kei Watanabe, Masaru Yagura, Yasukazu Nakamura

**Affiliations:** Kazusa DNA Research Institute, Kazusa-Kamatari, 2-6-7, Kisarazu, Chiba, Japan; Research and Development Section, Anicom Specialty Medical Institute Inc., 2-6-3 Chojamachi 5F, Yokohamashi-Nakaku, Kanagawaken, 231-0033 Japan; School of Life Sciences and the Center for Soybean Research of the State Key Laboratory of Agrobiotechnology, The Chinese University of Hong Kong, Shatin, Hong Kong SAR, China; National Institute of Genetics, Research Organization of Information and Systems, 1111 Yata, Mishima, Shizuoka, Japan

## Abstract

The domestic cat (*Felis catus*) is one of the most popular companion animals in the world. Comprehensive genomic resources will aid the development and application of veterinary medicine including to improve feline health, in particular, to enable precision medicine which is promising in human application. However, currently available cat genome assemblies were mostly built based on the Abyssinian cat breed which is highly inbred and has limited power in representing the vast diversity of the cat population. Moreover, the current reference assembly remains fragmented with sequences contained in thousands of scaffolds. We constructed a reference-grade chromosome-scale genome assembly of a domestic cat, *Felis catus* genome of American Shorthair breed, Anicom American shorthair 1.0 (AnAms1.0) with high contiguity (scaffold N50 > 120 Mb), by combining multiple advanced genomic technologies, including PacBio long-read sequencing as well as sequence scaffolding by long-range genomic information obtained from Hi-C and optical mapping data. Homology-based and *ab initio* gene annotation was performed with the Iso-Seq data. Analyzed data is be publicly accessible on Cats genome informatics (Cats-I, https://cat.annotation.jp/), a cat genome database established as a platform to facilitate the accumulation and sharing of genomic resources to improve veterinary care.

## Introduction

The domestic cat (*Felis catus*) is one of the most popular companion animals in the world. Originating in the Middle East, the domestication of cats began at least 10,000 years ago (Driscoll et al., 2007). Modern cat breeders have developed more than 40 breeds that are recognized by international cat registries, such as The Cat Fanciers’ Association (CFA) (Dennis-Bryan, 2013). Modern cat breeds have variations in traits including coat color and pattern, hair and tail length, and ear shape as well as genetic diseases (Bell et al., 2012). Such phenotypic variations among breeds are likely to have originate from genetic variations between breeds.

Since the release of the first draft genome of the domestic cat in 2007 (Pontius et al., 2007), studies on the genetic variations in *Felidae* have been accumulated rapidly. The population genetic structures were evaluated using microsatellite DNA markers (Lipinski et al., 2007; Menotti-Raymond et al., 2008; Kurushima et al., 2013) as well as genome-wide single-nucleotide polymorphisms (SNPs) (Gandolfi et al., 2018). These studies have also uncovered genetic variations among breeds. More recently, genome-wide analyses using high-throughput sequencers have identified breed-specific genetic variations and several candidate genes or genetic regions associated with diseases or traits (Aberdein et al., 2016; Lyons et al., 2016; Xu et al., 2016; Bertolini et al., 2016; Mauler et al., 2017; Buckley et al., 2020).

The currently available cat genome assembly Felis_Catus_9.0 (felCat9) is from the highly inbreeding Abyssinian cat (O’Brien et al., 2008). The genome has been much improved since its first release. However, there are still a large number of scaffolds. Variations captured using this reference genome could be limited due to the high inbreeding level, which may not well represent the general cat population. Such limitations potentially affect mapping accuracy and cause biases in variation calling and other genomic analyses. Therefore, for a deeper investigation into the genetic effects on the diseases and traits of cats, more high-quality cat genomes are essential.

Herein we established a high-quality chromosome-scale assembly named that we have named Anicom American Shorthair 1.0 (AnAms1.0) by integrating five high-throughput genomic technologies; (1) Pacific Biosciences (PacBio) Sequel deep sequencing, (2) error correction of the assembly by Illumina short-reads, (3) scaffolding by the chromatin conformation capture sequencing (Hi-C), and (4) optical mapping (OM) by the Bionano Saphyr system. The assembled genome sequences were aligned onto the felCat9 genome sequences for the construction of chromosome-level scaffolds. PacBio Iso-Seq full-length transcript sequencing data was employed for better gene-structure annotation. We further established a database, Cats-I, for the new cat genome assembly and its gene annotations, which is accessible at https://cat.annotation.jp.

## Materials and Methods

### Materials

A female American Shorthair cat, named Senzu (Fig. 1), was subjected to genome and transcriptome sequencing. We selected American Shorthair as the candidate breed for sequencing for the following three reasons. (1) The first one is that the breed is phylogenetically distant from and has a different genetic structure compared to the Abyssinian breed (Lipinski et al., 2007; Menotti-Raymond et al., 2008), i.e., the breed for which a high-quality genome assembly is already available (Buckley et al., 2020). (2) Several dozens of the genetic markers have uncovered American Shorthair cats has higher heterozygosity than that of Abyssinians (Lipinski et al., 2007), suggesting American Shorthair has more variations at the genome level. (3) The American Shorthair breed is one of the most popular cat breeds that can be found in most parts of the world (CFA 2020). Many other popular breeds such as the Scottish Fold breed originated from American Shorthairs, and the current breeding of such breeds allows outcrossing with American Shorthairs (Menotti-Raymond et al., 2008; CFA 2019). Together these factors make the American Shorthair breed is a good candidate for genome sequencing.

**Fig. 1.**
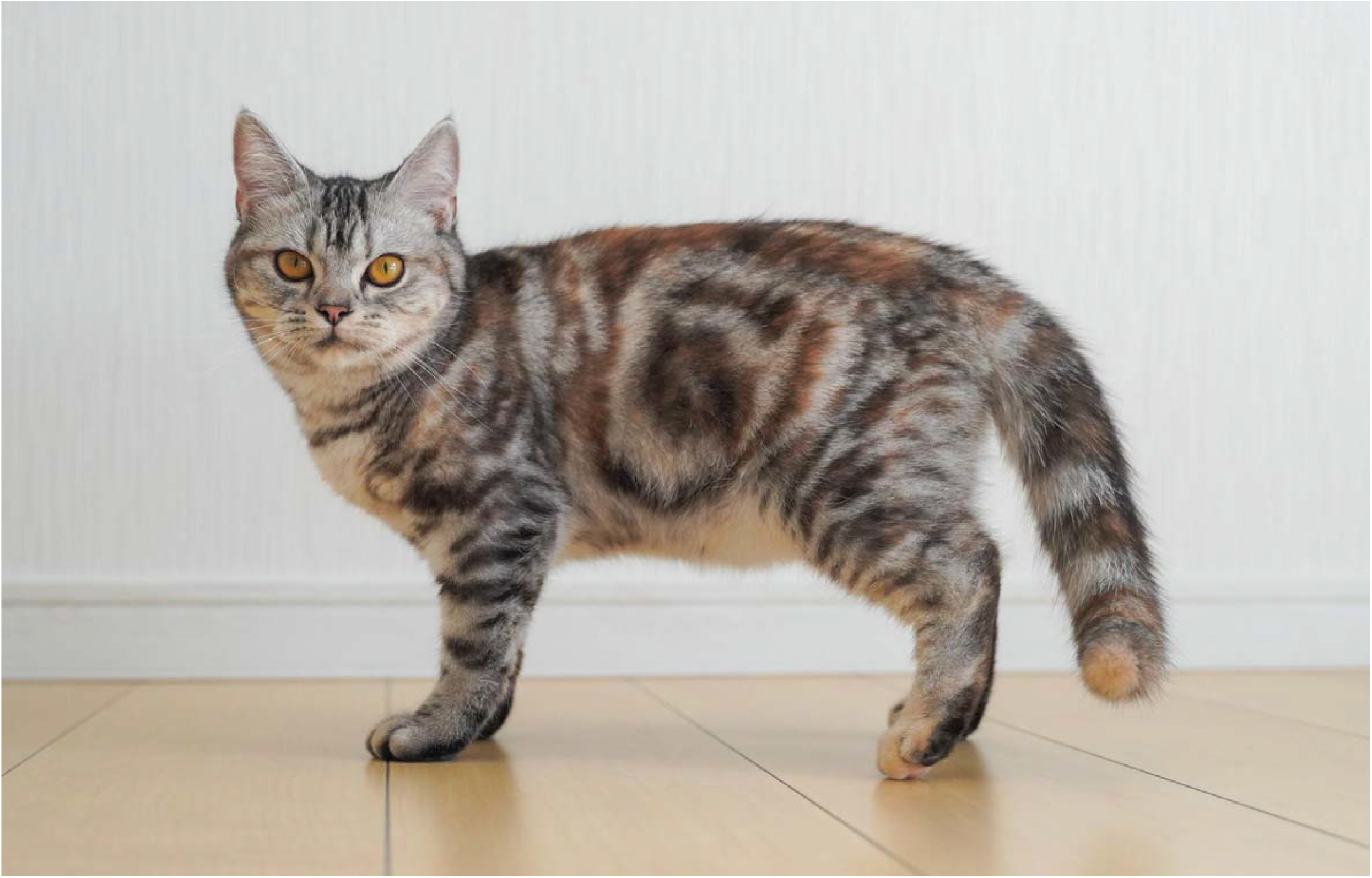
Photo of a female American Shorthair cat, named Senzu.

We selected the individual cat Senzu for sequencing by a pre-screening using an SNP genotyping array, the Infinium Feline 63K iSelect DNA array (Illumina, San Diego, CA), obtained from less invasive swab-based sampling and the blood. Fifteen American Shorthair cats were subjected to a genetic cluster evaluation based on the results of the ADMIXTURE analysis (Alexander et al. 2009) which included 40 cat breeds. Two cats, Senzu and Takae were shortlisted because the majority of their genetic components were derived from the American Shorthair breed. With consent from the owner, we collected blood, ovary, oviduct, and uterus samples of Senzu and Takae during contraceptive surgery, and used them for the subsequent experiments. We selected Senzu for genome assembly construction because of her more inbred genetic background compared to Takae. Samples collected from Takae were used for the gene prediction and annotation by Iso-Seq.

### Illumina and PacBio sequencing

The Genomic DNA of Senzu was extracted from her blood with the use of a DNeasy Blood and Tissue Kit (QIAGEN, Germantown, MD, USA). Sequence libraries were prepared using the SMRTbell Express Template Prep Kit 2.0 (PacBio, Menlo Park, CA, USA). The size selection of the library was performed by BluePippin (Sage Science, Beverly, MA) to remove DNA fragments less than 30 kb in length. The library was then sequenced using a Sequel system (PacBio) with 28 SMRT cells. The sequence reads were assembled using FALCON v.1.8.8 (c) with the length cutoff of 34 kb, in order to generate primary contig sequences and to associate contigs representing alternative alleles. Haplotype - resolved assemblies, i.e. primary contigs and haplotigs, were generated using FALCON_Unzip ver. 1.8.8 (Chin et al., 2016). Potential errors in the resultant primary contig sequences were corrected twice using ARROW ver. 2.2.1 implemented in SMRT Link v.5.0 (PacBio). Paired-end (PE) libraries were constructed for the Senzu’s DNA and sequenced using an Illumina HiSeqX system (Illumina, San Diego, CA) with the read-length of 151 bp. The obtained short reads were used for further error-correction of the contig sequences by the tool Pilon (Walker et al. 2014). The short read data were also used for genome size estimation using Jellyfish ver. 2.1.4 (Marcais and Kingsford, 2011).

### Hi-C scaffolding

A Hi-C library was constructed from an ovary from Senzu by using a Dovetail™ Hi-C Kit with *DPNII* restriction enzyme (Dovetail, Scotts Valley, CA, USA). The library was sequenced by the Illumina HiSeqX to generate 150 bp PE reads. The obtained sequence reads were then aligned onto the PacBio contigs for scaffolding using the HiRise pipeline (Putnam et al., 2016).

### Bionano optical mapping and Hybrid Assembly

High molecular weight genomic DNA was extracted from cat uterus by following the Bionano Prep Animal Tissue DNA Isolation Kit instructions (Bionano, San Diego, CA) using the Soft Tissue Protocol (Document no. 30077 Rev C). Optical mapping data were then generated on the Bionano Saphyr platform following the Bionano Direct Label and Stain (DLS) protocol (i.e., Document no. 30206 Rev F). Briefly, flash-frozen tissue was homogenized and embedded into agarose gel to prevent mechanical shearing during nuclear DNA isolation. DNA was then recovered from the molten gel by drop dialysis, followed by labeling with DLE-1 enzyme to produce sequence motif-specific fluorescent signal patterns and by YOYO-1 backbone staining to visualize the molecule lengths. Fluorescence-labelled single molecules were stretched and had images captured automatically in nanochannel arrays in Saphyr Chip G1.2 (Part no. 20319) on the Bionano Saphyr system (Part no. 60325). Optical maps containing molecule length and label distance information were converted from images by Bionano Access ver. 1.4. We performed the d*e novo* map assembly with Bionano Solve ver.3.4.1 and Bionano Tools ver. 1.4.1 via Bionano Access v1.4 using non-human, no preassembly, non-haplotype-aware, no SV mask, and cut complex multi-path region (CMPR) parameters. We also performed the hybrid scaffolding of Pilon-polished Hi-C scaffold sequences with Bionano Solve ver. 3.4.1 and Bionano Tools ver. 1.4.1 via Bionano Access v1.4 setting in order to resolve conflicts in both the optical map and sequence assemblies.

### Construction of chromosome-level scaffolds

Chromosome-level scaffolds were constructed by aligning scaffolds onto the felCat9 genome (https://www.ncbi.nlm.nih.gov/assembly/GCF_000181335.3/) by RagOO (Alonge et al. 2019). The completeness of the assembly was assessed by BUSCO ver. 3 with the mammalian *odb9* gene sets (Simão et al., 2015). The genome comparison was performed by the Nucmer module in MUMmer 3.23 (Kurtz et al. 2004). Known repetitive sequences registered in Repbase (https://www.girinst.org/repbase/) and *de novo* repetitive sequences defined by RepeatModeler 1.0.11. (http://www.repeatmasker.org/RepeatModeler) were identified by RepeatMasker 4.0.7. (http://www.repeatmasker.org/RMDownload.html).

### Transcriptome analysis

Total RNAs were extracted from ovary, oviduct, and uterus of two American Shorthair cats, Senzu and Takae. The RNA extracted from multiple organs were mixed to prepare an Iso-Seq library in accordance with the manufacturer’s protocol (PacBio). The library was sequenced by a Sequel system with two SMRT cells. The obtained reads were clustered using the Iso-Seq 3 pipeline implemented in SMRT Link ver.8.0 (PacBio) and then mapped onto the assembled genome with Minimap2 (Li et al. 2018), and collapsed to obtain nonredundant isoform sequences using a module in Cupcake ToFU (https://github.com/Magdoll/cDNA_Cupcake).

### Gene prediction and annotation

The empirical gene predictions were performed as follows. First, the coding sequences (CDS, Felis_catus.Felis_catus_9.0.cds.all.fa) and cDNA (Felis_catus.Felis_catus_9.0.cdna.all.fa) sequences of felCat9 genome and the Iso-Seq sequence fragments in our data (Ensembl ver. 98.9; Felis_catus_9.0-based) were mapped to the assembled genome using GMAP (ver. 2019-06-10, Wu et al., 2005). *Ab initio* gene predictions were also performed by BRAKER (v2.1.0, Hoff et al., 2016, 2019; Stanke et al., 2006, 2008) with the AUGUSTUS *ab initio* option. The empirical and *ab initio* gene prediction results were combined to form final gene predictions. Differences between the CDS of felCat9 and the Iso-seq fragments mapped regions, as well as differences in genes predicted by BRAKER, were used as novel gene candidates. We performed the following: the regions where the CDS of felCat9 was mapped was replaced with N, and the CDS was extracted using the gff (the file extension) made by mapping the Iso-Seq fragments to the unmasked genome. Similarly, the regions which were mapped both the CDS of felCat9 and the Iso-Seq fragments were replaced with N. The CDS regions were extracted from the N-masked genome using the gff (the file extension) predicted by BRAKER. We considered the CDS regions without N as novel gene candidates. The extracted CDS from both the CDS of felCat9 -mapped regions and the novel gene candidates were translated into protein. The functional annotation was performed using InterProScan (v5.33-72.0, Jones et al., 2014), and the KEGG Automatic Annotation Server (KAAS, Moriya et al., 2004), and RPS-BLAST (Marchler-Bauer and Bryant, 2004).

### Base and structure variances against Felis_catus_9.0

Base variances (SNPs and Indels) and structural variants (SVs) of the assembled genome were identified against the felCat9 genome. For the detection of base variants, Illumina PE reads of Senzu were mapped onto both the assembled genome and the felCat9 genome by Bowtie2 (Langmead and Salzberg, 2012), and variant calls were performed by bcftools 1.9 in SAMtools (Li et al. 2009). Possible false base variants were filtered out with the following conditions: exclude multi alleles, minimum DP = 20, maximum DP = 100 minimum GQ = 50, minimum QUAL = 200, max-missing = 1.0. SVs were identified by using the pbsv module in SMRT Link 6.0 (PacBio).

## Results and Discussion

### Genome assembly construction

A total of 1.54 Gb reads were obtained from an Illumina PE library. The distribution of distinct k-mers (k=17) shows two peaks at multiplicities of 77 and 154, which project an estimated genome size of 2.44 Gb and 1.22 Gb respectively (Fig. 2). According to Pontius et al. (2007), the estimated genome size of a domestic cat is 2.7 Gb. Our present estimated genome size of 2.44 Gb is less than that, but closely similar to the total length of the felCat9 genome. It is thus predicted that the peak of multiplicities of 77 was generated from homozygous regions of the genome of Senzu. The absence of a peak derived from heterozygous regions of the two genomes suggests highly homozygosity of the Senzu genome.

**Fig. 2.**
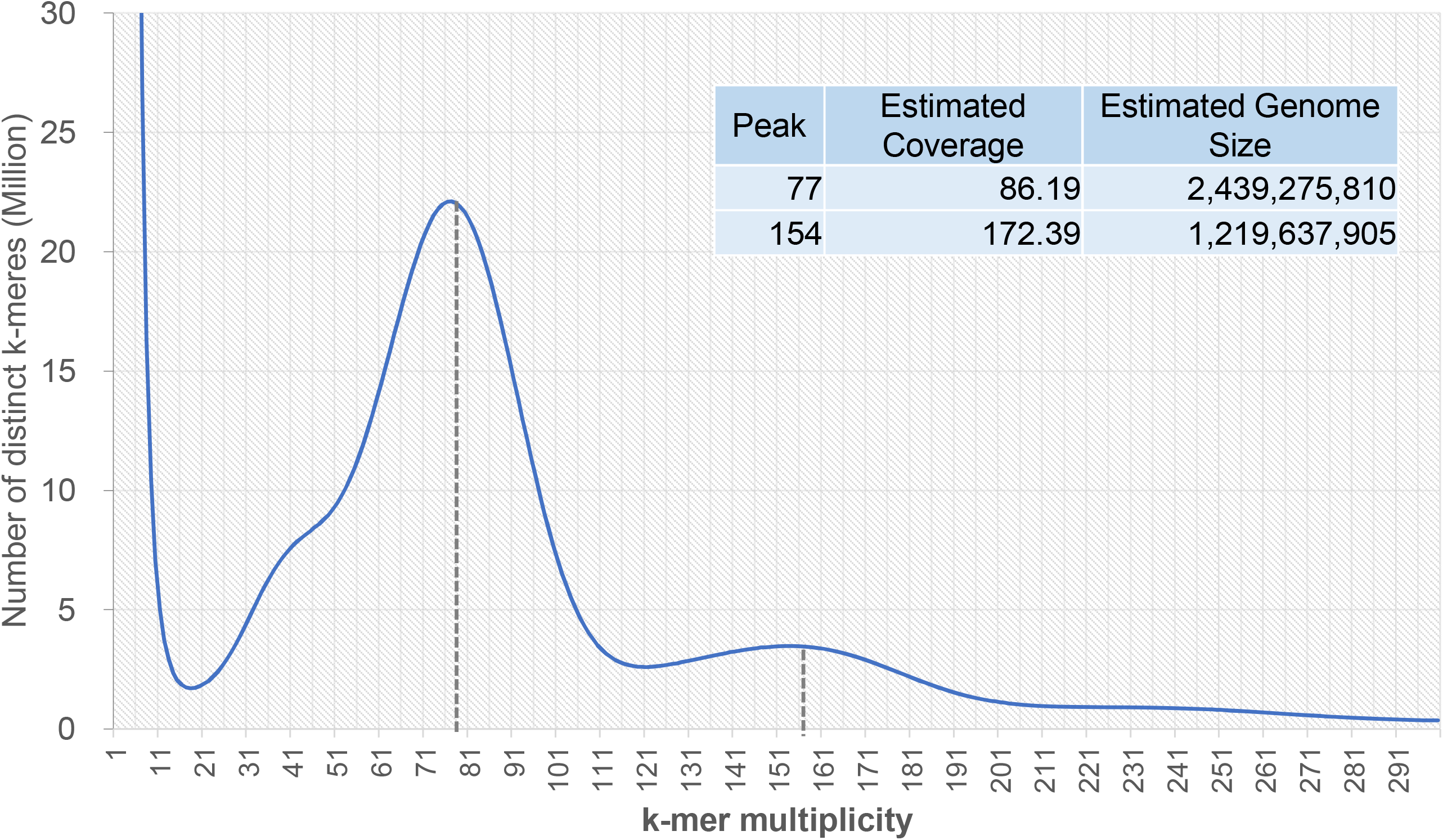
Distribution of the number of distinct k-mar (k=17) of an American Shorthair cat, Senzu, with the given multiplicity values.

### PacBio assembly

A total length of 345.4 Gb of PacBio reads was generated from 28 SMRT cells and read lengths ≥ 35 kb were used for assembly. The total length and coverage of the Senzu genome (2.44Gb) were 120.7 Gb and 49.5x, respectively. The Falcon unzip assembly generated 547 primary and 5,512 haplotig contigs. The total length of the primary contigs was 2.47 Gb with the N50 length of 21.3 Mb. The genome sequences of the primary contigs were scaffolded with N100 by Hi-C reads after the error collection with PacBio and Illumina reads. Table 2 provides the numbers of generated scaffolds, with the total and N50 lengths of 2.47 Gb and 149.6 Mb, respectively. The total N length of the Hi-C scaffolds was 17,500 bp.

**Table 1.** Status of genome assembly with Pacbio, Illumina PE and Hi-C reads.

|  | FALCON unzip contigs |  | Hi-C scaffolds |
| --- | --- | --- | --- |
|  | Primary | Haplotigs |  |
| Number of sequences | 547 | 5,512 | 372 |
| Total length (bp) | 2,468,876,196 | 1,611,610,682 | 2,472,554,877 |
| Average length (bp) | 4,513,485 | 292,382 | 6,646,653 |
| Max length (bp) | 75,142,039 | 4,345,118 | 241,446,035 |
| Min length (bp) | 19,724 | 109 | 19,548 |
| N50 length (bp) | 21,265,058 | 621,279 | 149,647,023 |
| A | 717,851,700 | 470,824,785 | 718,272,105 |
| T | 717,392,275 | 470,630,412 | 719,070,471 |
| G | 516,692,290 | 335,012,532 | 517,690,152 |
| C | 516,939,931 | 335,142,953 | 517,504,649 |
| N | 0 | 0 | 17,500 |
| Total (ATGC) | 2,468,876,196 | 1,611,610,682 | 2,472,537,377 |
| GC% (ATGC) | 41.9 | 41.6 | 41.9 |

**Table 2.** Status of genome assembly after optical mapping (OM) hybrid scaffolding.

|  | Hi-C scaffolds | Post-OM hybrid scaffolding (total) | Post-OM hybrid scaffolding (Super-Scaffolds) |
| --- | --- | --- | --- |
| Number of sequences | 372 | 348 | 37 |
| Total length (bp) | 2,472,554,877 | 2,493,108,843 | 2,458,290,041 |
| Average length (bp) | 6,646,653 | 7,164,106 | 66,440,271 |
| Max length (bp) | 241,446,035 | 109,076,552 | 222,017,658 |
| Min length (bp) | 19,548 | 1,381 | 100,606 |
| N50 length (bp) | 149,647,023 | 109,076,552 | 109,076,552 |
| A | 718,272,105 | 719,036,553 | 709,861,598 |
| T | 719,070,471 | 718,318,909 | 709,089,249 |
| G | 517,690,152 | 517,486,211 | 509,273,155 |
| C | 517,504,649 | 517,715,032 | 509,513,899 |
| N | 17,500 | 20,552,140 | 20,552,140 |
| Total (ATGC) | 2,472,537,377 | 2,472,556,703 | 2,437,737,901 |
| GC% (ATGC) | 41.9 | 41.5 | 41.4 |

### Optical mapping

Optical maps provide whole-genome long-range genomic information, generated from single-molecule data from over a hundred-kilobase pairs up to megabase pairs long, to resolve large genomic structures. A total of 1,225,034 optical mapping (OM) molecules over 150 kb labeled with DLE-1 enzyme recognition patterns were obtained from the Bionano Saphyr run, with N50 length of 336,375 bp, containing 383.6 Gb (Fig. 3A). The molecules were assembled into 118 genome maps containing 2,419 Mb with a chromosome-level N50 length of 73.1 Mb (Fig. 3B).

**Fig. 3.**
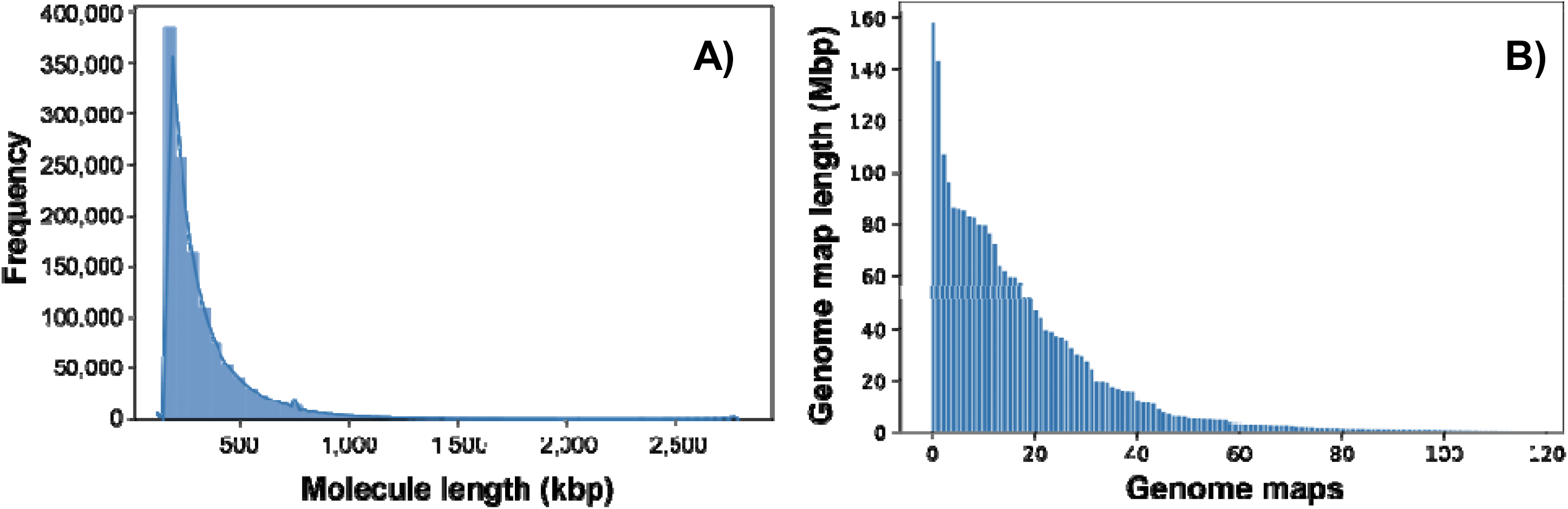
Length distribution of optical mapping data A) Length distribution of raw molecules from the OM data. B) Length distribution of assembled genome maps the OM mapping data

Among the 372 input sequences, 87 were joined into 37 super-scaffolds containing 2.46 Gb, i.e. 99.4% of the input sequences (Table 2 and Fig. 4A). The changes in non-ambiguous bases were resulted from the insertion of the DLE-1 sequence motif at the fluorescent label signals and the trimming of the sequences at the scaffolding and conflict resolution, whereas the increase in N-bases indicates the sizing of gaps to facilitate further sequence placement. Decreases in the scaffold N50 and maximum length were the results of the resolution of assembly conflicts, indicating the correction of mis-assemblies (Fig. 4B).

**Fig. 4.**
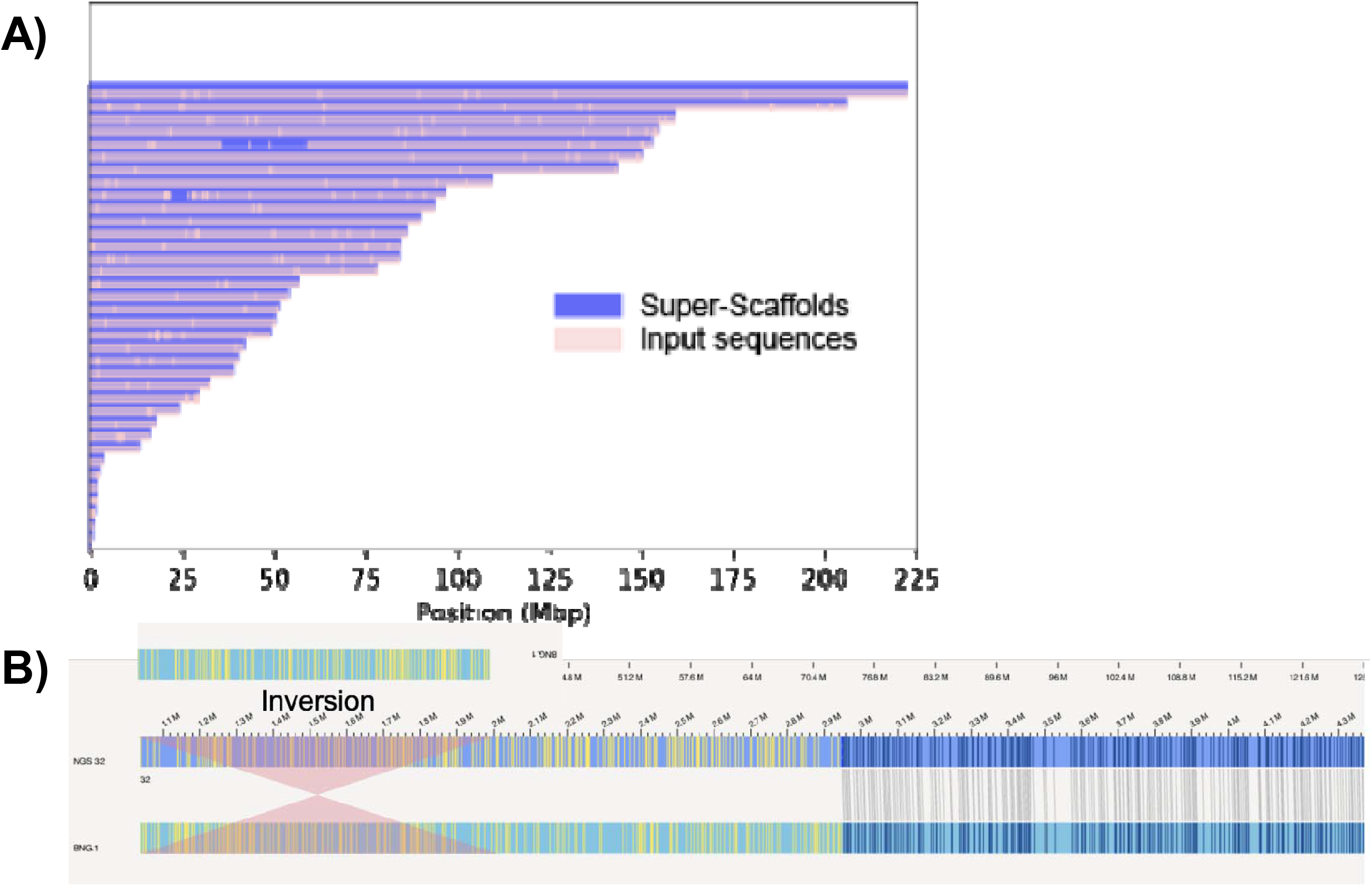
Results of optical mapping hybrid scaffolding A) Visualization of input Hi-C sequences scaffolded by OM data into 37 super-scaffolds. B) Example of misassembly correction by OM hybrid scaffolding.

### Chromosome-level scaffolding

We aligned the scaffolds sequences modified by optical mapping onto the felCat9 genome by RagOO and designated the genome as ‘AnAms1.0’. The statistics of the AnAms1.0 and felCat9 genomes are provided in Table 3. AnAms1.0 consists of 19 chromosome-level scaffolds and only one short unmapped scaffold, the length of which was 1,381 bp. The 19 chromosome-level scaffolds represent 18 autosomal chromosomes and one sex (X) chromosome. The corresponding chromosome numbers are given according to the felCat9 genome. The total length of AnAms1.0 is 2,493,141,643 bp, and the N50 is 151.1 Mb. The lengths of the 19 chromosome-level scaffolds in AnAms1.0 range from 41.75 Mb (E3) to 243.50 Mb (A1). The comparison of the total ATGC length of the 19 scaffolds between the two genomes, revealed that AnAms1.0 is 57,627,227 bp longer than felCat9. We investigated the assembly quality of AnAms1.0 by mapping the sequences onto 4,104 BUSCOs (i.e., benchmarking universal single-copy orthologs), and the results demonstrated that the number of complete BUSCOs was 3,923 (95.6%), including 3,907 (95.2%) singlecopy genes and 16 (0.4%) duplicated genes. The numbers of fragmented and missing BUSCOs were 99 and 82, respectively.

**Table 3.** Statistics of the AnAms1.0 and felcat genomes.

|  | AnAms1.0 |  | felCat9 |  |
| --- | --- | --- | --- | --- |
|  | All scaffolds | Scaffolds correspond to nucleic chromosomes | All scaffolds | Scaffolds correspond to nucleic chromosomes |
| No. of sequences | 20 | 19 | 4,508 | 19 |
| Total length, bp | 2,493,141,643 | 2,493,140,262 | 2,521,863,845 | 2,460,251,910 |
| Avg. length, bp | 124,657,082 | 131,217,909 | 559,420 | 129,486,943 |
| Max. length, bp | 243,504,654 | 243,504,654 | 242,100,913 | 242,100,913 |
| Min. length, bp | 1,381 | 41,750,578 | 1,969 | 44,648,284 |
| N50 length, bp | 151,107,676 | 151,107,676 | 149,751,809 | 149,751,809 |
| A | 718,322,616 | 718,322,255 | 721,302,099 | 703,154,254 |
| T | 719,032,844 | 719,032,444 | 721,735,169 | 703,748,672 |
| G | 517,632,441 | 517,632,143 | 516,811,973 | 504,031,439 |
| C | 517,568,802 | 517,568,480 | 516,603,929 | 503,993,730 |
| N | 0.83 | 0.83 | 1.8 | 1.84 |
| Total, ATGC | 2,472,556,703 | 2,472,555,322 | 2,476,453,170 | 2,414,928,095 |
| GC%, ATGC | 41.9 | 41.9 | 41.7 | 41.7 |

A graphical view of the syntenic relationships between AnAms1.0 and felCat9 for the 19 scaffolds is shown in Figure 5. The alignment of homologous sequence pairs along each scaffolds shows high similarity between the two genomes, with the exception of the B2 chromosome.

**Fig. 5.**
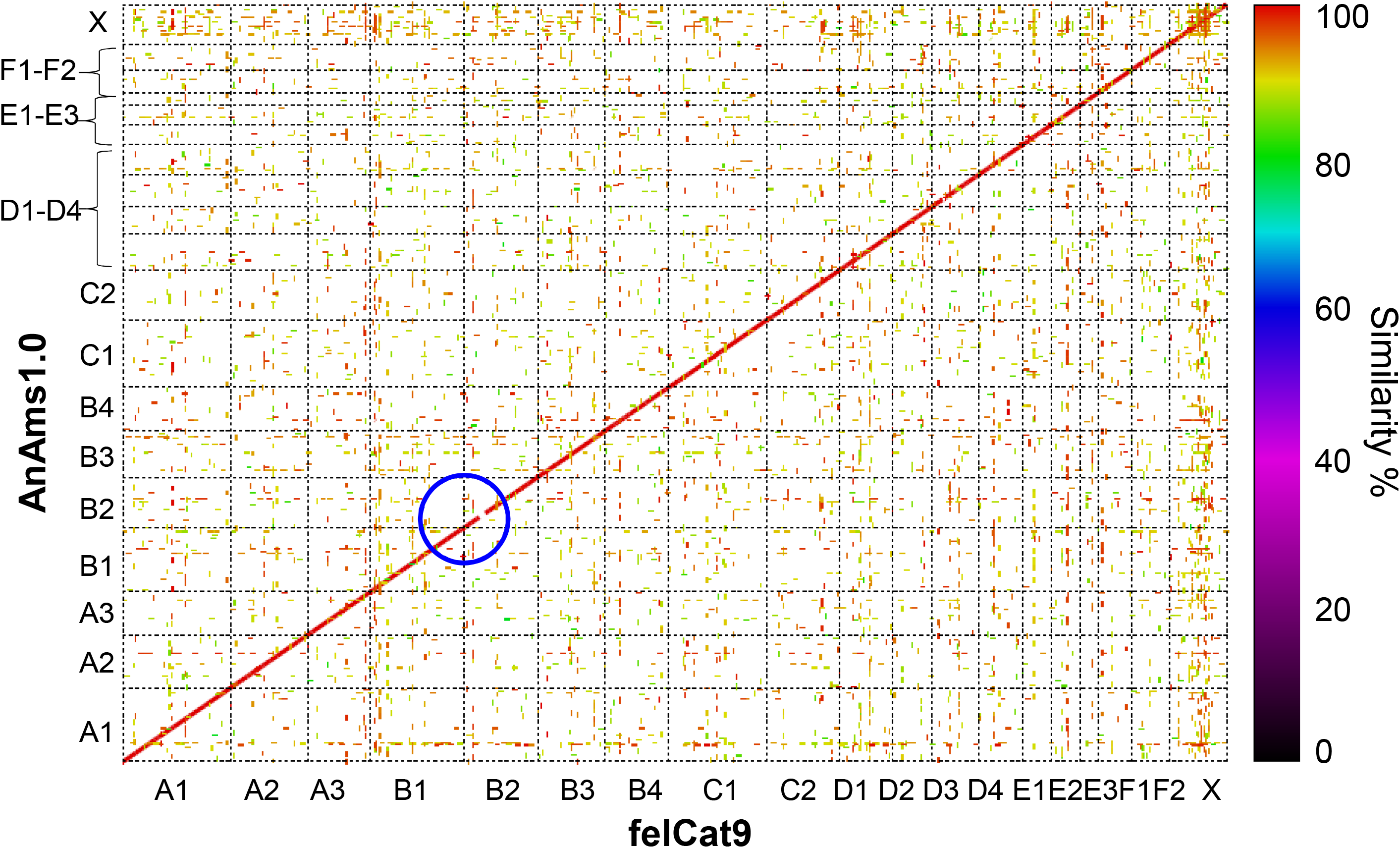
Genome sequence comparison of the 19 chromosomes of AnAms1.0 against that of felCat9. A blue circle shows incompatible regions on B2 chromosome.

### Repetitive sequences

The total length of repetitive sequences in the AnAms1.0 genome was 880.2 Mb, occupying 35.3% of the assembled genome (Table 4). The ratio of repeat sequences was similar to that for the felCat9 (34.6%). The percentages of repetitive sequences on each chromosome-level scaffold ranged from 32.6% (B2) to 36.6% (B3) in the autosomal chromosomes of AnAms1.0, and that of the X chromosome was 49.1% (Figure 6).

**Table 4.**
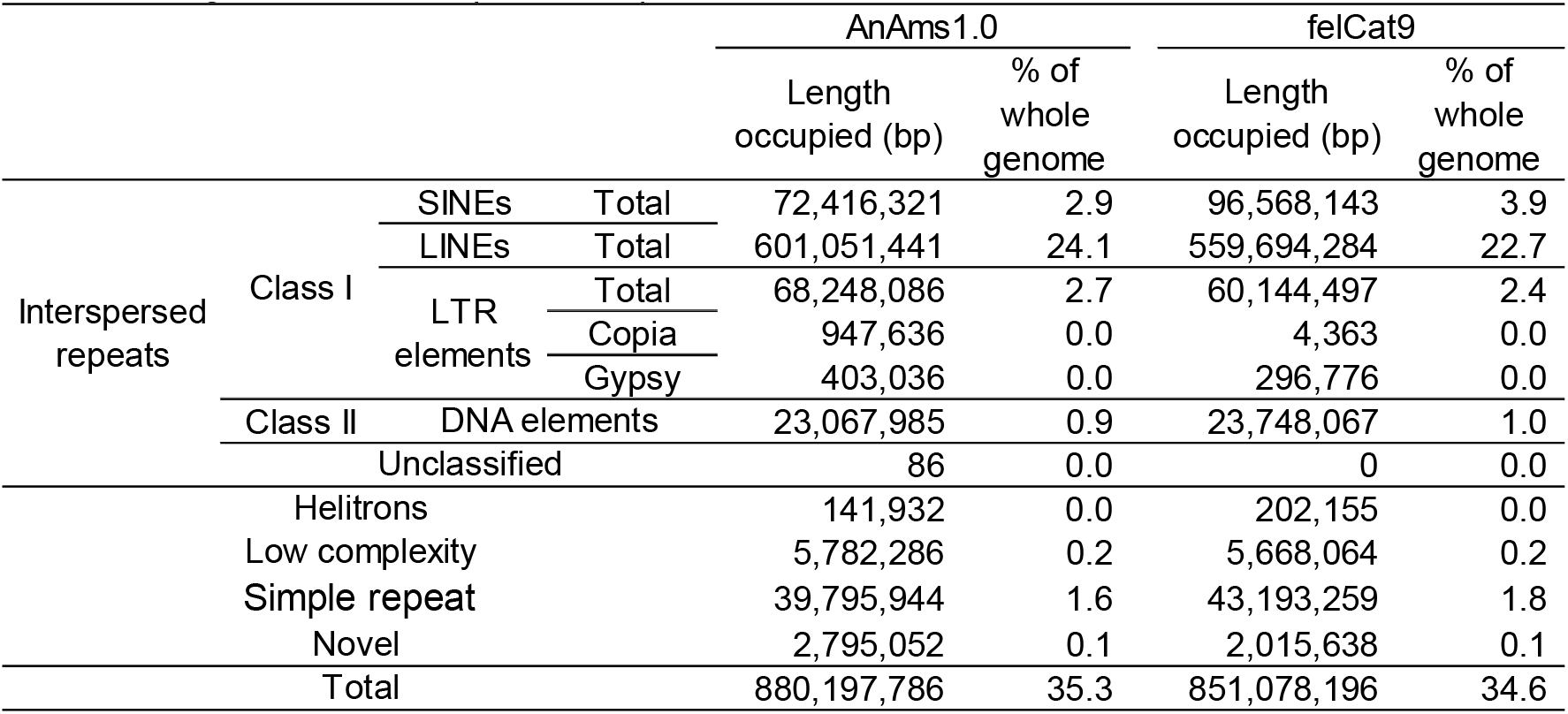
Length and ratio of repetitive sequences.

**Fig. 6.**
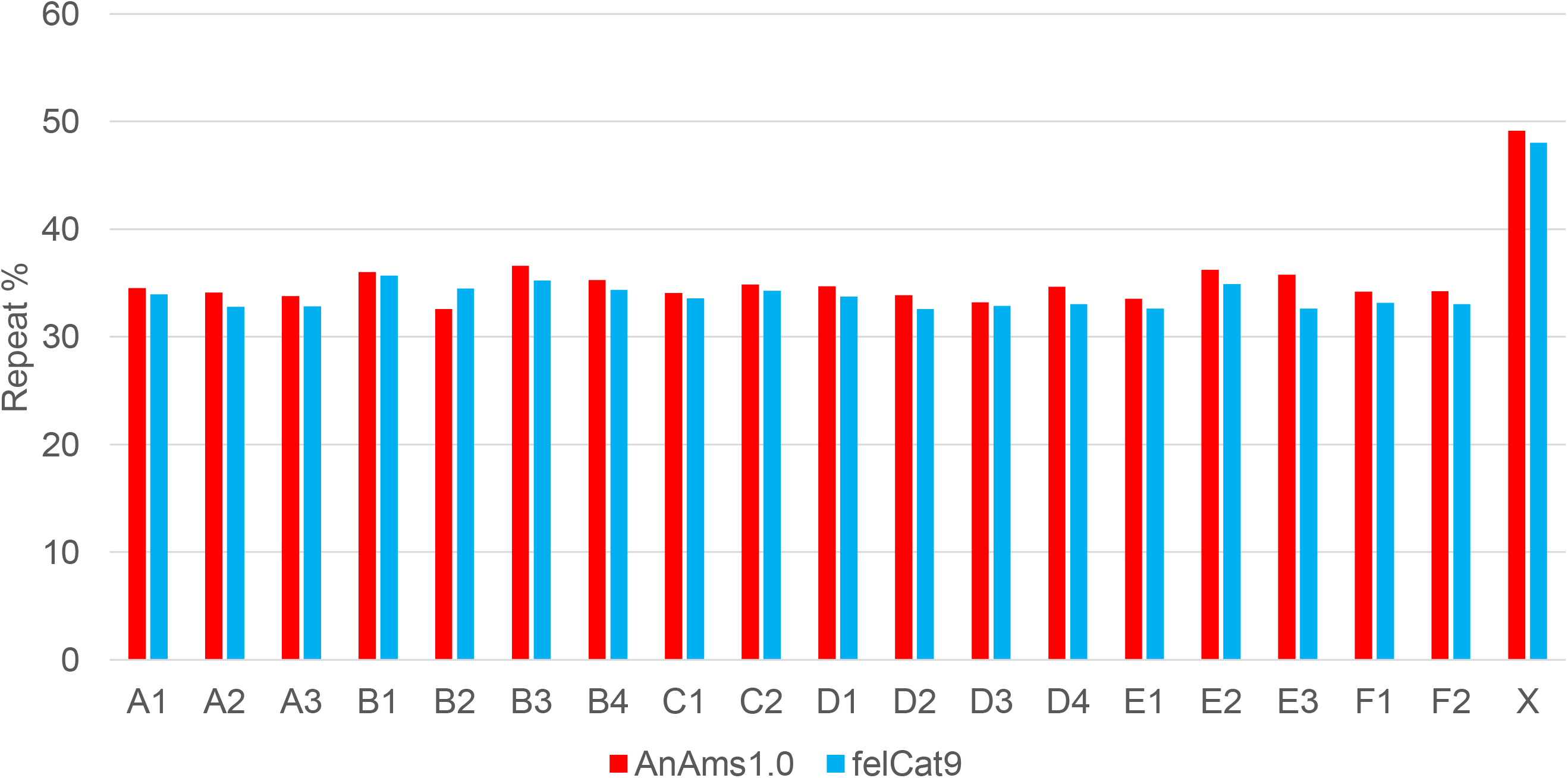
Percentages of repetitive sequences on each chromosome-scale scaffolds

### Iso-Seq analysis

Iso-Seq sequences totaling 20.9 Gb and 31.6 Gb in length were obtained from the Senzu and Takae samples respectively. The sequences from the two individual cats were integrated and clustered by Iso-Seq3, and the high-quality (hq) sequences were collapsed and filtered based on quality by Cupcake ToFU. As shown in Table 5, during the process of assembly, the total number of resultant sequences decreased from 50,436 (Iso-Seq3/hq) to 26,662 (collapsed/filtered). However, the ratio of complete BUSCOs did not differ greatly among the IsoSeq3/hq, collapsed, and collapsed/filtered results. The ratios of complete BUSCOs ranged from 46.7% to 47.1%, and that of missing BUSCOs ranged from 40.8% to 41.3%. In the Iso-Seq analysis, transcript sequences were obtained from ovary, oviduct, and uterus. We suspect that this sampling of a limited number of organs is a probable cause of the large portion of missing BUSCOs.

**Table 5.** Statistics of Iso-Seq assembly from ovary, oviduct, and uterus of Senzu and Takae.

|  | Iso-Seq3/hq | collapsed | collapsed/filtered |
| --- | --- | --- | --- |
| Number of sequences | 50,436 | 38,150 | 26,662 |
| Total length (bp) | 80,707,453 | 61,698,768 | 44,915,792 |
| Average length (bp) | 1,600 | 1,617 | 1,685 |
| Max length (bp) | 8,077 | 8,077 | 8,077 |
| Min length (bp) | 55 | 84 | 85 |
| N50 length (bp) | 1,761 | 1,789 | 1,859 |
| BUSCOs % (n=4,104) |  |  |  |
| Complete | 47.1 | 46.8 | 46.7 |
| Complete single | 25.9 | 29.5 | 31.5 |
| Complete double | 21.2 | 17.3 | 12.5 |
| Fragmentized | 12.1 | 12 | 12 |
| Missing | 40.8 | 41.2 | 41.3 |

### Gene prediction and annotation

Of the 19,587 protein-coding genes of the felCat9 genome, 19,562 genes were mapped on AnAms1.0; of the 26,662 sequences obtained with Iso-Seq, 26,614 sequences were mapped on AnAms1.0; the mapping results of felCat9 transcripts and Iso-Seq sequences identified 23,199 loci on AnAms1.0. Protein domain annotations were obtained for the translation products of these loci by InterProScan, KAAS, and RPS-BLAST. The gene models predicted by BRAKER with the *ab initio* option are available in our genome browser.

### Base and structure variances against felCat9

Illumina PE reads were mapped onto the AnAms1.0 and felCat9 genomes for the detection of base variances (SNPs and indels). The numbers of identified candidate variants on AnAms1.0 and felCat9 genomes were 4,124,481 and 8,688,925 respectively. The variants of the repetitive sequences were filtered out and the remaining numbers of variants on the AnAms1.0 and felCat9 genomes were 2,544,090 and 5,550,296, respectively. Of the 2,554,090 variants identified in AnAms1.0, 2,652,448 were heterozygous (hetero), and 1,642 were alternative homozygous (alt homo). It was considered that the 1,642 alt homo variants were generated by mis-calling or mis-assembly of the AnAms1.0.

The numbers of hetero and alt homo variants identified on the felCat9 genome are 2,731,328 and 2,818,968, respectively. The density of heterozygous variants identified onto the two genomes shows similar values, ranged from 95 variants / 1 Mb (E3) to 1,228 variants / 1Mb (C1) in AnAms1.0 and from 99 variants / 1 Mb (E3) to 1,242 variants / 1Mb (C1) in felCat9.0 (Figure 7). The result suggested that the bias of heterozygosity between the two haploid Senzu genomes. The density of alt homo variants identified on felCat9 ranged from 232 variants / 1 Mb (X) to 1,217 variants / 1Mb (A1). The alt homo variants reflect the sequence differences between AnAms1.0 and felCat9 genomes. The results suggest that sequences of the sex chromosome were more conserved than autosomal chromosomes.

**Fig. 7.**
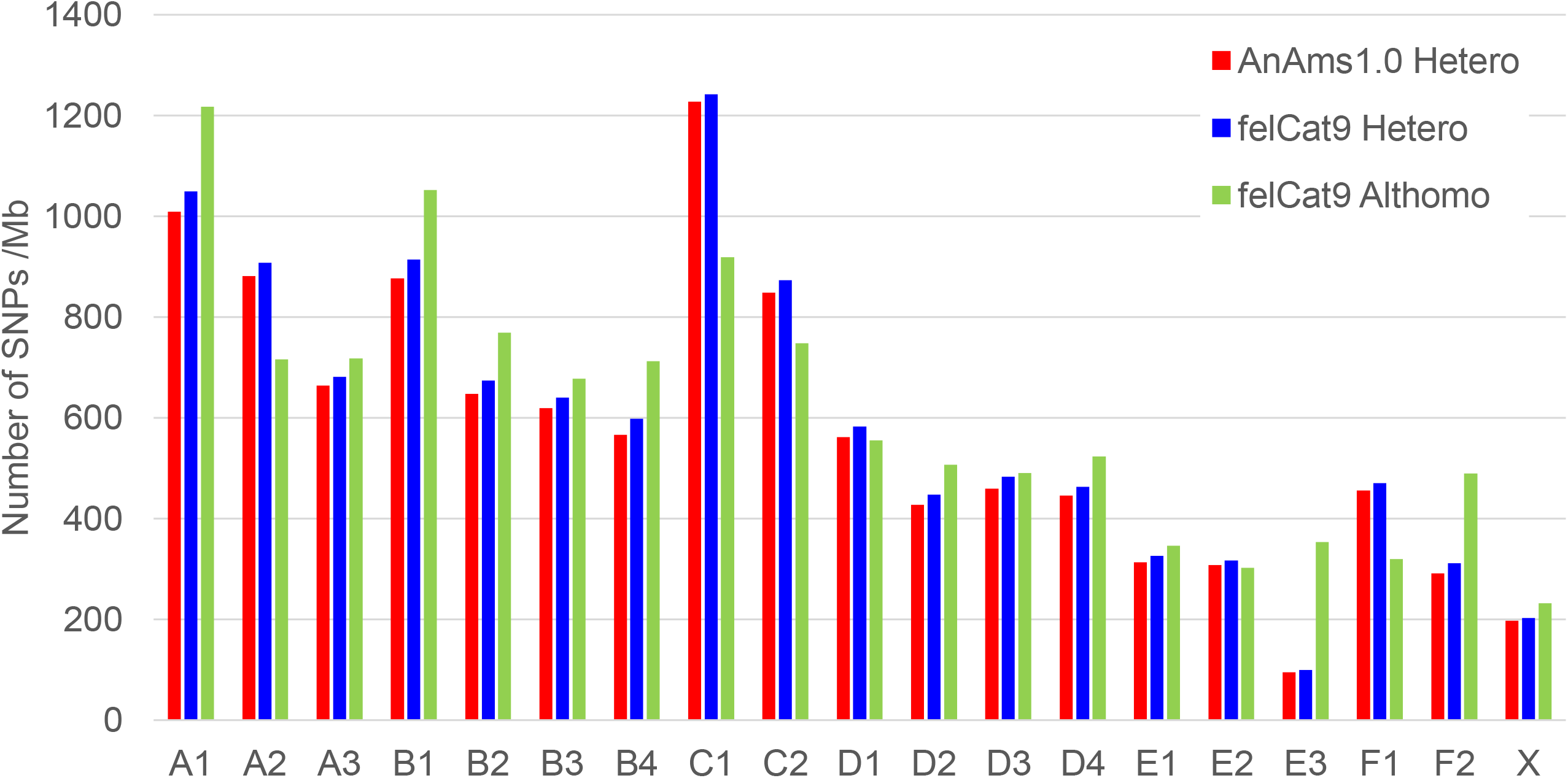
Average density base variants identified by mapping Senzu Illumina reads onto each chromosome of AnAms1.0 and felcat9.

In order to identify SVs, we mapped Senzu PacBio reads onto the felCat9 genome. A total of 16,035,301 PacBio sub reads were mapped across the felCat9 genome with the mean coverage of 120.1x. The number of identified SVs was 155,535 with the total length of 45,222,456 bp (Table 6). The frequently observed SVs were deletions (55.1%) and insertions (39.3%).

**Table 6.**
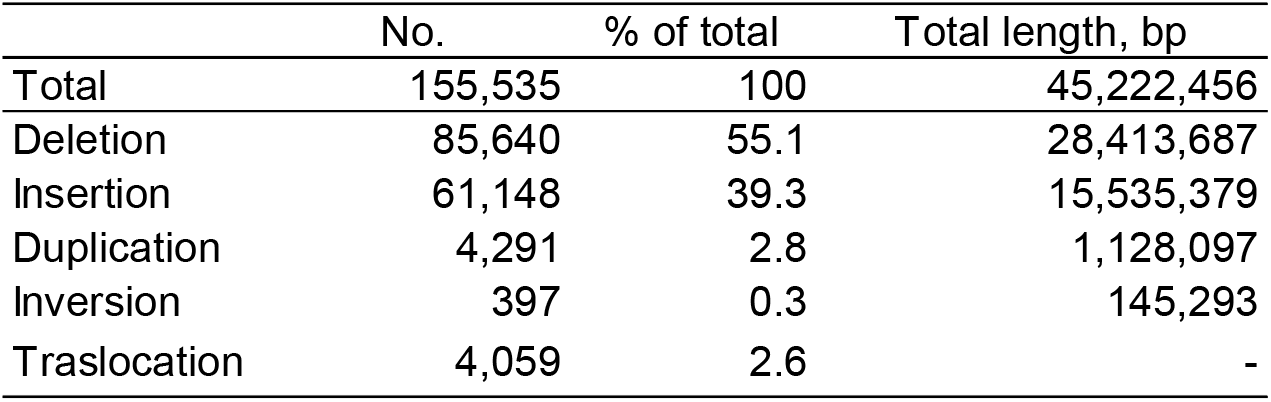
Number of SVs of the Senzu genome against felCat9.

The distribution of the mapped PacBio read depth shows large insertion and deletion regions of the Senzu genome against felCat9 (Figure 8). Possible insertions were observed on most of the chromosomes. The large deletion was identified on E3 chromosome. The large deletion identified on the B2 chromosome by Nucmer was not detected in the SV analysis.

**Fig. 8.**
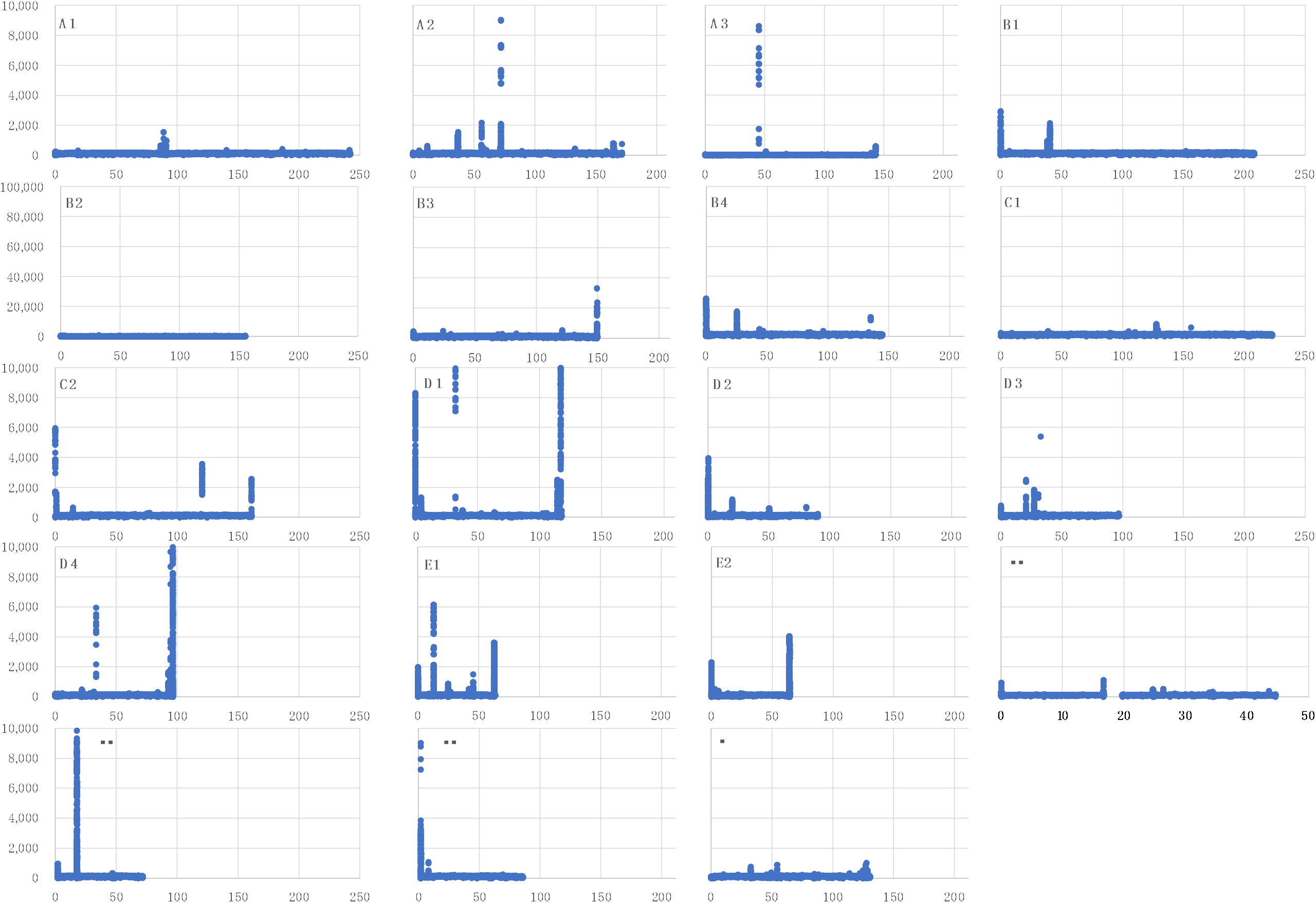
Distribution of mapped Senzu PacBio read depth on the felCat9 genome. X and axis are physical position (Mb) and mapped read depth (x), respectively. The scale of E3 is changed from other graphs to show a deletion of the chromosome.

### Database construction

Cats-I, a genome database for AnAms1.0 containing a genome browser with annotated genes, expression profiles, BLAST search tools, and download tools for genomic resources are available at https://cat.annotation.jp

### Implications for veterinary medicine and precision medicine in cat

The developed genome assembly, AnAms1.0., potentially provides insight into veterinary medicine as well as precision medicine in the domestic cat. We found over 60 thousand novel SVs in the AnAms1.0. against to felCat9. The SV has the potential to affect inherited diseases or severe phenotypes (Buckley et al., 2020). Although further genomic analyses to detect such variants are required, one of the advantages of AnAms1.0. has the potential to map more variants compared to felCat9.

Precision medicine is an advanced medical treatment concept which chooses the optimum preventive medical treatment and therapy that takes into account individual differences in genetic characteristics, environment, and lifestyles specific to the patient. In the U.S.A., the individualized medical treatment research by large-scale individual genome information acquisition and epidemiological study of the living environment is promoted on the development of the cancer therapy in recent years by President Obama’s Precision Medicine Initiative (https://obamawhitehouse.archives.gov/the-press-office/2015/01/30/fact-sheet-president-obama-s-precision-medicine-initiative). In human medicine, this approach has been made possible by the preparation of the high-quality human reference genome as a foundation and the large-scale accumulation and integration of disease-related gene annotation and genetic variation information such as Clinvar (Landrum et al. 2013). Recent studies have shown in precision medicine in companion animals (Mauler et al., 2017, Mealey et al., 2019). Here, we have constructed a chromosome-level genome assembly for a breed of cat, and are about to launch an information foundation for pet precision medicine for the future of the veterinary care.

## Data Availability

The assembled genome sequences have been submitted to the DDBJ/ENA/NCBI public sequence databases under the BioProject ID PRJDB9879 with the accession numbers AP023152-AP023171.

## Acknowledgments

This work was supported by the Research Foundation of the Foundation of Anicom Specialty Medical Institute Inc, Kazusa DNA Research Institute, the Hong Kong Research Grants Council Collaborative Research Fund (C4057-18EF), the CUHK Group Research Scheme 3110135 to TFC and Hong Kong Research Grants Council Area of Excellence Scheme (AoE/M - 403/16) to HML and TFC. Computations were partially performed on the NIG supercomputer at the ROIS National Institute of Genetics.

